# Vitamin D deficiency alters prostate epithelial differentiation and increases prostate cancer aggressiveness in ex vivo and in vivo models

**DOI:** 10.64898/2026.02.16.700850

**Authors:** Adriana Duraki, Kirsten D. Krieger, Sasha Celada, Robert A. Holt, Ryan M. Brown, Luyu Wang, Michael J. Schlicht, Maarten C. Bosland, Robert M. Sargis, Donald Vander Griend, Larisa Nonn

**Author notes:** Corresponding Author: Larisa Nonn, PhD, Professor of Pathology, 840 S Wood St, Room 130 CSN, MC 847, Chicago, IL 60612 USA. **DISCLOSURE STATEMENT:** The views expressed in this article are those of the authors and do not necessarily reflect the position or policy of the Department of Veterans Affairs or the United States government. R.M.S. has received honoraria from CVS/Health unrelated to this manuscript. The other authors declare no competing interests. **DATA ACCESS STATEMENT:** Research data supporting this publication are available at NCBI GEO GSE304873, GSE309716, and GSE311628.

## Abstract

Here, we examined the consequences of biologically relevant vitamin D deficiency, a known risk factor for aggressive prostate cancer, using ex vivo and in vivo models. Phenotypic and single-cell RNA sequencing of mouse prostate organoids showed that vitamin D deficiency stunted luminal cell differentiation more than androgen deficiency, which is a known driver of prostate development. Mice fed a vitamin D-deficient diet showed significantly altered expression of androgen-responsive genes in their prostate luminal cells, as determined by single-cell RNA sequencing. MDA-PCa-2b human prostate cancer cells, when maintained for 6 months in 1,25-dihydroxyvitamin D, lost the ability to form xenografts, despite normal proliferation in vitro. RNA sequencing showed that these cells also had disruptions in androgen signaling and multiple cancer-related pathways. This study offers new insights and validation of vitamin D’s role in both benign and malignant prostate biology, underscoring its essential hormonal functions and supporting strategies for vitamin D supplementation to reduce prostate cancer risk in vulnerable populations.

**STATEMENT OF SIGNIFICANCE:** Vitamin D is an essential hormone, however, the non-calcemic consequences of vitamin D deficiency remain poorly defined, despite its high prevalence in the population. This study demonstrates significant biological consequences of vitamin D deficiency on prostate cells at biologically relevant levels in multiple systems.

## INTRODUCTION

Prostate cancer (PCa) is the second most common cancer in men worldwide, with over 1.46 million cases reported in 2022.^1^ PCa often progresses slowly and in high-income countries it has a favorable prognosis, with 5-year survival rates of 70–100%.^2^ PCa shows the greatest racial disparity among cancers in the U.S., with non-Hispanic Black men facing higher risks of diagnosis (17% vs. 12.7%) and mortality (3.1% vs. 2.1%) compared to non-Hispanic White men.^3^ This disparity likely reflects a complex interplay of socioeconomic factors, healthcare access, life-style factors, environmental exposures, and genetic/biological differences, particularly variations in androgen receptor signaling and other genomic alterations.^4,5^

Vitamin D deficiency is prevalent in the U.S. and elsewhere and has been linked to PCa aggressiveness via in vitro, in vivo, and population studies. The vitamin D receptor (VDR) is highly expressed in prostate tissue, and treatment of human PCa cell lines and primary epithelial cells with the active metabolite, 1,25-dihydroxyvitamin D (1,25D), results in growth suppression, suggesting a role in prostate homeostasis.^6,7^ Because vitamin D is synthesized in the skin by UV radiation, factors that reduce vitamin D synthesis, such as age, race, dark skin color, and northern latitude, may increase the risk of PCa.^8^ Vitamin D deficiency is associated with increased all-cause cancer mortality in men.^9^ Geographic and seasonal associations between PCa mortality and prognosis exist, further suggesting a link between vitamin D and PCa.^10,11^ Vitamin D status appears to have a stronger association with aggressive or lethal disease^12–14^ than with overall PCa incidence.^15–17^ Vitamin D pathway genetic variants are also associated with recurrence and lethal PCa.^18^ There is also evidence to suggest that vitamin D supplementation benefits other prostate conditions, including benign prostate hyperplasia (BPH) and lower urinary tract symptoms (LUTS), with clinical trials reporting reduced prostate growth, PSA levels, and symptom severity.^19–21^

Studies using mouse models have consistently demonstrated that vitamin D status alters PCa progression. Dietary studies in TgAPT_121_ and TRAMP mice have shown that vitamin D deficiency accelerates proliferation, high-grade Prostatic Intraepithelial Neoplasia formation, tumor invasion, and metastasis.^22–24^ Prostate-specific VDR deletion in the TgAPT_121_ model resulted in increased epithelial proliferation, more and larger PCa foci, indicating that loss of VDR signaling promotes tumor development.^25^ A vitamin D-enriched diet reduced the growth of PC-3 xenograft tumors in mice compared to standard diets.^26^ Collectively, these models demonstrate that systemic or genetic disruption of vitamin D pathways enhances PCa initiation and progression.

In the U.S., vitamin D deficiency (<20 ng/mL) affects 29% of adults, with the highest prevalence among non-Hispanic Blacks (72%) compared to Hispanics (43%) and non-Hispanic Whites (19%).^27^ Given that Black men experience both higher PCa mortality and greater vitamin D deficiency, numerous studies have examined this link. In prospective studies, low 25D levels were associated with increased PCa risk and aggressiveness in African American men.^28–30^ Siddappa et al. identified a distinct VDR cistrome in African American PCa.^31^ These data underscore the need for research to clarify the role of vitamin D in PCa progression, including in vitamin D-deficient and high-risk groups.

Here, we aimed to gain a deeper understanding of the protective effects of vitamin D on the prostate by using biologically relevant models of vitamin D deficiency. Ex vivo prostate development, in vivo dietary studies, and experiments with PCa cell lines revealed profound actions of vitamin D on prostate cells and PCa biology.

## MATERIALS AND METHODS

### Mouse Colony/Maintenance

All animals were housed in the pathogen-free barrier facility managed by the University of Illinois at Chicago (UIC) Animal Care and Use Program, which is an AALAC-accredited facility. The facility complies with all National Institutes of Health standards for the care and use of vertebrate animals. All mouse work protocols were reviewed and approved by the Office of Animal Care and Institutional Biosafety (OACIB) under Animal Care Committee (ACC) protocol numbers 22-079 (C57BL/6 mice and vitamin D diet study) and 21-018 (xenograft mice). All surgeries were completed in designated surgery rooms located within the COMRB Animal Facility corridor.

### Mouse Prostate digestion to single cells

C57BL/6 male mice were euthanized, the prostates removed, and the AP, DL, and V lobes were dissected.^53,54^ Tissues were dissociated into single cells using an optimized protocol adapted from Drost et al..^55^ Briefly, tissues were macerated using a sterile scalpel into < 1 mm^3^ pieces, then digested at 37°C while rotating for 30-45 minutes in 5 mg/mL collagenase type II (Gibco 17101015), 10 mM nicotinamide (Sigma N0636), 10 μM Y-27632 (Sigma Y0503), 1 nM DHT, 1 U/uL DNase I (STEMCELL 07470), and Advanced Dulbecco’s Modified Eagle Medium/Ham’s F-12 (AdvDMEM/F-12; Gibco 12634028). Digestion was stopped by adding base media containing 10 mM nicotinamide, 10 μM Y-27632, 1 nM DHT, and AdvDMEM/F-12, and the tissues were pelleted. Following one minute treatment with 1 mM EDTA (Invitrogen AM9260G) tissues were dissociated by vigorous pipetting with a pre-cut p1000 pipette tip for 2 minutes in 1 U/mL Dispase (STEMCELL 07923). Single cells were isolated through a 30-micron cell filter (Miltenyi Biotec 130-041-407) four times. Cell suspensions were then transferred to DNA LoBind Tubes (Eppendorf 022431021) and centrifuged at 300 x g for 3 minutes. Pellets were gently washed in 1% Bovine Serum Albumin (BSA; Jackson Immunoresearch 001-000-162). Cells were counted using a hemocytometer.

### Organoid Culture

Primary whole prostate or lobe-specific prostate cells were isolated as described above. Organoid culture methods were adapted and optimized from our previously published human organoid methods^56^ and mouse organoid methods.^55^ Briefly, primary dissociated cells were counted and prepared in 33% Matrigel Matrix (Corning 356234) combined with media (100 μL/well) at either 10,000 cells/well (freshly dissociated cells) or 25,000 cells/well (previously frozen dissociated cells). The media contained 1 μM Y-27632, 1 nM DHT, 5% Charcoal Stripped Fetal Bovine Serum (CS-FBS; Corning 35-072-CF), and AdvDMEM/F-12. Cells were plated ultra-low attachment 96 well plates (Corning 3474) and incubated at 37°C. Fresh media was added every two to three days. The treatment media contained 1,25D (Enzo BML-DM200-0050) at various concentrations, as specified in the figure legends.

### Organoid Imaging and Phenotypic Analysis

All imaging was performed using an EVOS FL Auto 2 Cell Imaging System (Thermo). Throughout the experiments, 4X and 10X bright-field images were taken to assess individual organoid morphology, while whole-well acquisition imaging (WWAI) was completed to assess whole-well differences between conditions. The size and shape of each individual organoid were quantified using Celleste Image Analysis Software as previously described by our group.^56^ Organoids were also classified as either “Luminal” or “Other” within Celleste software, allowing for analysis of specific visual morphologies. Organoid relative area (size) data were exported and graphed using Prism (GraphPad).

### Murine Prostate Tissue Gene Expression

Following euthanasia, prostates were removed, microdissected into individual paired lobes (anterior, dorsolateral, and ventral), and flash frozen immediately in liquid nitrogen. Tissues were pulverized in a tissue homogenizer with TRIzol reagent (Invitrogen 15596026). Homogenized samples in TRIzol were then stored at -80 °C. Kidney tissue was also collected and processed in the same manner as a positive control.

### RNA Isolation, cDNA Synthesis, and Gene Expression

RNA was isolated according to the TRIzol manufacturer’s instructions and quantified using a Nanodrop OneC (Fisher Scientific). cDNA was synthesized from 500 ng RNA using the High-Capacity cDNA Reverse Transcription Kit (Applied Biosystems 4368814). qPCR was run with FastStart Universal SYBR Green Master (Rox; Roche) on the QuantStudio 6 Real-Time PCR System (Thermo). Gene expression results were analyzed using the ddCT method. Expression of Vdr, Ar, and Cyp24a1 was normalized to TATA-binding protein (Tbp) housekeeper gene (**Table S4**).

### Organoid Passaging

At 14 days of culture, DL prostate organoids were digested into single cells, counted, and re-seeded at 1000 cells per well in 3D culture as organoids as previously described. Organoids were imaged and analyzed on day 14, as previously described.

### Organoid Culture and Dissociation for Single-Cell RNA Sequencing

Primary dorsolateral prostate cells from WT mice were dissociated and plated as organoids as described above. The treatment media for this experiment consisted of four conditions: vehicle control, 10 nM DHT, 25 nM 1,25D, and 10 nM DHT combined with 25 nM 1,25D. 14 day organoids were dissociated for scRNAseq. Briefly, the growth media were removed, and dispase was added, followed by incubation at 37 °C for 40 minutes. Cells were then mechanically dissociated using a pre-cut pipette tip, and collected (5 minutes at 300 x g). The pellet was resuspended and washed in 10% Fetal Bovine Serum (FBS; Gibco 26140079) in PBS (Corning 21-040-CV). Cells were incubated in TrypLE (Gibco 12605010) for 20 minutes at 37 °C, followed by washes. Cells were run through 40-micron cell strainer (Flowmi BAH136800040). Cell quantification and viability assessment, determined by the Trypan Blue exclusion assay, were performed using the Countess Automated Cell Counter (Fisher). 10000 cells per sample were used.

### Single Cell RNA Sequencing of Organoids

All samples exhibited >85% cell viability prior to processing with the 10X Genomics Chromium Next GEM Single Cell 3’ workflow (CG000204 Rev D). Libraries were prepared using the Chromium Next Gem Single Cell 3’ GEM, Library and Gel Bead Kit v3.1 (PN-1000128), Chromium Next GEM Chip G Single Cell Kit (PN-1000127), and Chromium i7 Sample Index Plate (PN-220103), with a targeted recovery of 5,000 cells per sample. Libraries were sequenced on one SP lane with 2×90nt reads in the NovaSeq 6000 (Illumina) at the University of Illinois at Urbana-Champaign (UIUC) DNA Services. Sequencing depth was targeted at 50,000 reads per cell. Initial alignment and quality control were performed using the aggr and Cell Ranger pipelines at the UIUC Biotechnology Center, with subsequent alignment to the NCBI murine reference genome (GRCm39). Detailed scRNAseq quality metrics are provided in **FIGURES S1-S2** and **TABLE S2.**

Downstream analysis was conducted using Seurat (v4.3.0). Cells were excluded if they expressed fewer than 2000 genes, exhibited high mitochondrial gene expression (>10%), or were doublets. SCTransform (Seurat; glmGamPoi method) was used to normalize, select highly variable genes, and transform the data using the top 3000 variable genes with both mitochondrial contamination (mitoRatio) and cell cycle differences (CC.Difference) regressed out. Principal component analysis (PCA) followed by uniform manifold approximation and projection (UMAP) dimensionality reductions were computed using the top 20 principal components, with clustering resolutions of 0.4 for non-integrated data and 0.2 resolutions for integrated data. Seurat’s integration analysis was also performed.^57^ For integrated analyses, PCA/UMAP/TSNE embeddings and cluster assignments from the integrated dataset were transferred to the non-integrated data to preserve biological variability that could be lost during integration. Data are available in NCBI GEO GSE304873.

### Sequencing Cluster Identification and Marker Analysis

Cluster-specific markers were identified by differential expression analysis comparing each cluster to all other cells in the dataset using Seurat’s FindAllMarkers function with default parameters (non-parametric Wilcoxon rank sum test). Conserved markers were identified using Seurat’s default FindConservedMarkers function to identify genes that are consistently expressed across all samples within a given cluster. Differentially expressed genes (DEGs) between specified sample groups and individual clusters were identified using Seurat’s FindMarkers function. Clusters were assigned cell-type identities based on established marker genes reported in published prostate scRNAseq datasets from both murine and human studies.^58–62^ Mouse orthologs of various markers were derived from human cell type markers for use in our murine dataset using the NCBI Gene database.^63^ All scRNAseq plots including VlnPlots, FeaturePlots, ggplots, PCA plots, UMAP plots, heatmaps, and DotPlots were generated using Seurat in R.

### Gene Set Enrichment Analysis

The Hallmark Collection gene set (Mus musculus species) from the Molecular Signatures Database (MSigDB) was obtained using the R package msigdbr (v7.5.1). Identified DEGS were ranked by average log^2^ fold change, and Gene Set Enrichment Analysis (GSEA) was performed utilizing the R package fgsea (v1.24.0) and visualized using the plotGseaTable function.^64^

### Vitamin D-Deficiency Mouse Diet Study

8-week-old male C57BL/6 mice (Jackson Laboratory, Bar Harbor, ME USA) were randomly assigned to cages and diets. Mice were fed ad libitum either the vitamin D-control diet (2.2 IU/g; 2200 IU/kg; Teklad Custom Diet #TD.89124) or the vitamin D-deficient diet (0.05 IU/g; 50 IU/kg; Teklad Custom Diet #TD.89123). Sample size calculations were based on previously observed differences in vitamin-D-treated organoids, indicating that n = 3 mice per group provided 80% power to detect a ≥2-fold difference (SD ≤0.4) using an IACUC-approved calculator (Boston University). The study included n = 6 mice per diet, distributed across the following endpoints: scRNAseq (n = 3 per diet) or formalin fixation and histology (n = 3 per diet).

Animals were monitored weekly for overall health and body weight. Humane endpoints were predefined in accordance with the AVMA and UIC OACIB. No adverse health events or severe weight loss occurred during the study. Mice were euthanized after 24 weeks on the diets.

### Serum Vitamin D Levels (LC-MS/MS)

Blood samples were collected at euthanasia via cardiac puncture, allowed to coagulate at room temperature for 1h, and centrifuged at 1500 x g for 15 min at 4°C. Serum was flash-frozen and stored at -80°C until analysis. Serum 25-hydroxyvitamin D (25D) levels were quantified using a validated LC-MS/MS assay as previously described^65^ by Heartland Assays (Ames, IA).

### Histology

Whole prostates were dissected, fixed in 10% neutral-buffered formalin (Sigma HT501128), and subsequently transferred to 70% ethanol. The anterior prostate lobe was placed into a separate cassette, while the remaining prostatic lobes were maintained en bloc with the urethra and bladder ring attached to maintain orientation during sectioning. Tissues Processing, paraffin embedding, and sectioning (5 µm) were performed by the Research Histology and Tissue Imaging Core at UIC. H&E images were examined by a veterinary pathologist (M.C.B.).

### Single Cell RNA Sequencing of Vitamin D Diet Prostate Tissue

Dorsolateral prostate lobes were dissected, cleared of surrounding adipose tissue, and dissociated into single-cell suspensions as previously described. Final suspensions were passed through a 40-micron cell strainer (Flowmi BAH136800040) and assessed using trypan blue exclusion (NanoEntek EBT-001) with an automated cell counter (Countess, Thermo) and verified using a manual hemocytometer. All samples contained >75% viable cells, with three of four samples exhibiting >90% viability. Libraries were prepared using a dual index strategy following the 10X Genomics Chromium Next GEM Single Cell 3’ workflow (CG000315 Rev E), using the Library and Gel Bead Kit v3.1 (PN-1000268), Chromium Next GEM Chip G Single Cell Kit (PN-1000127), and Dual Index Kit TT Set A (PN-1000215), with targeted recovery of 5,000 cells per sample. Dual index libraries were sequenced on two S4 lanes with 2×150nt reads in the Novaseq 6000 (Illumina) at the UIUC DNA Services facility, targeting a sequencing depth of 100,000 reads per cell. Initial alignment and quality control were performed using aggr and Cell Ranger pipelines at the UIUC Biotechnology Center, with alignment to the NCBI murine genome (GRCm39). Detailed scRNAseq quality metrics are provided in **FIGURE S3** and **TABLE S3**.

Downstream analyses were conducted using Seurat (v4.3.0) with initial exclusion as previously described. SCTransform (Seurat; glmGamPoi method) was utilized for data normalization, identification and selection of highly variable genes, and data transformation, employing the top 3000 variable genes with regression of both mitochondrial contamination (mitoRatio) and cell cycle differences (CC.Difference). PCA was performed, followed by UMAP using the top 15 principal components. Clustering resolution was set to 0.1 for non-integrated analyses and 0.05 for integrated data analysis. Data integration was performed as described previously, with integrated PCA/UMAP/TSNE embeddings and cluster assignments transferred back to the non-integrated datasets. Cell-type annotation, differentially gene expression analyses, and data visualization were performed as previously described. Data are available on NCBI GEO GSE309716.

### Extended Growth of Prostate Cancer Cells in Vitamin D

MDA-PCa-2b cells were used for all PCa cell line experiments. Cells were obtained from ATCC and authenticated by short-tandem repeat (STR) profiling at the University of Illinois at Urbana-Champaign Cancer Center. Mycoplasma testing was performed using PCR-based detection (Sigma MP0035) at UIC Genome Research Core (GRC). MDA-PCa-2b cells were grown on plates/flasks coated with FNC Coating Mix (AthenaES 0407) in BRFF-HPC-1 media (AthenaES 0403) supplemented with 20% FBS. Cells were maintained in culture at 37°C with 5% CO_2_ and passaged 1:3 once cells reached 70-80% confluence.

Following the establishment of consistent growth, MDA-PCa-2b cells were expanded and divided into three parallel culture conditions containing vehicle control, 1 nM, or 10 nM 1,25D. Cells were maintained under these conditions for six months. Treatment media were prepared fresh at each change to reduce hormone degradation and were replaced every 3-4 days.

### MDA-PCa-2b Xenografts

Castrated 8-week-old male SCID mice (Envigo; C.B-17/IcrHsol-Prkdc-scid) were used for the PCa xenograft experiments. To standardize the circulating hormone levels, all mice were castrated and supplied with androgen through a testosterone-containing subcutenous silastic implant. Cells were xenografted two weeks after castration and testosterone implantation. The three vitamin D-adapted MDA-PCa-2b cell lines (Vehicle, 1 nM 1,25D, and 10 nM 1,25D) were injected in 75% Matrigel Basement Membrane Matrix (LDEV-free; Corning 354234) and 25% Hanks’ Balanced Salt Solution (HBSS; Gibco 14170-112) at 1 × 10^7^ cells/mL. Each mouse received two 100uL subcutaneous injections (1 million cells/injection, one on each flank (2 grafts/mouse; 10 grafts/cell treatment). After tumors became palpable, caliper measurements were performed biweekly, increasing to three times per week once any tumor reached 500 mm^3^. Mice were euthanized at a tumor volume of 1000 mm³ or upon meeting a humane endpoint criterion. Tumors were excised, weighed, and photographed.

### In vitro Growth Assays

The MDA-PCa-2b cells of the three vitamin D-6 month treatments (vehicle, 1 nM 1,25D, and 10 nM 1,25D) were counted using the Countess Automated Cell Counter and diluted in base media (20% FBS BRFF-HPC-1) to 3.0 × 10^5^ cells/mL. Cells (15,000 cells/well) were seeded in FNC-coated 96-well plates and incubated in the Incucyte S3 (Sartorius) for live imaging. Vehicle 6-month cells were used for 4-day treatment with 1 and 10 nM 1,25D. Phase images were acquired every 4 hrs for 72hrs using the Adherent Cell-by-Cell module on the Incucyte (Sartorius, US). Analysis excluded objects < 300 um^2^ to avoid quantifying detached/dead cells. Phase area confluence ratios normalized to 0d0h0m, average phase object area (um^2^) data, and images from each condition and timepoint were exported for graphing using Prism.

### Short-term Treatment vs. 6 Month 1,25D Gene and Protein Expression

The MDA-PCa-2b cells of the three vitamin D-6 month treatments (Vehicle, 1 nM 1,25D, and 10 nM 1,25D) were plated in (20% FBS BRFF-HPC-1) (15,000 cells/well) 2.0 × 10^6^ cells/well in FNC-coated 6-well plates. Vehicle 6-month cells were used for 24h or 48h treatment with 1 and 10 nM 1,25D. RNA isolation was performed as described above for VDR, CYP24A1, AR, KLK3 and FKBP5 with normalization to TBP (**Table S4**).

### Western Blot Analysis

Media was aspirated and cells were washed with ice-cold 1X HEPES Buffered Saline and lysed in 1.5X Cell Lysis Buffer (Cell Signaling Technology (CST) 9803), supplemented with Protease/Phosphatase Inhibitor Cocktail (CST 5872) and Phenylmethanesulfonyl Fluoride (CST 8553). Plates were incubated on ice for 10 mins, centrifuged at 12,000 rpm for 15 mins, and supernatants were collected. Protein concentration was determined using a Bradford assay (BIO-RAD 5000006) on a Nanodrop OneC. Samples were prepared at 1 μg/μL in NuPAGE LDS Sample Buffer (Invitrogen NP0007), NuPAGE Sample Reducing Agent (Invitrogen NP0009), and diH_2_O, then heated to 70°C for 10 minutes. 30 µg of protein were separated on 4-12% Bis-Tris gel (NuPAGE; Invitrogen NP0335BOX) using MOPS SDS running buffer (Invitrogen NP0001) and transferred to a PVDF membrane (Invitrogen PB5210) using the Power Blotter XL System (Invitrogen PB0013). Membranes were blocked (LI-COR 927-60001) for 1hr at room temperature and incubated overnight at 4° C with primary antibodies specific to VDR (D2K6W; CST 12550), AR (PG-21; EMD Millipore 06-680), FKBP5 (D5G2; CST 12210), and housekeeper H3 (96C10; CST 3638). All primary antibodies were diluted 1:1000 in blocking buffer with 0.2% Tween 20 (BIO-RAD 1706531). Following washing in 1X Tris Buffered Saline (TBS) with 0.1% Tween 20 (TBS-T), membranes were probed with respective secondary antibodies depending on species of primary antibody, including IRDye® 800CW Goat anti-Mouse IgG Secondary Antibody (LI-COR 926-32210) and IRDye® 680RD Goat anti-Rabbit IgG Secondary Antibody (LI-COR 926-68071). Secondary antibodies were diluted 1:1000 and incubated for 1hr at room temperature. Blots were imaged using the Odyssey CLx Imager (LI-COR). Band intensity was quantified using the Image Studio Software (LI-COR; Version 5.2.5). Relative protein expression results were analyzed by comparing the protein of interest expression to the histone H3 and normalized to vehicle -treated controls.

### Total RNA Sequencing

Total RNA was isolated from triplicate cultures of 24 h and 6 month 1,25D treated MDA-PCa-2b cells using the RNeasy Plus Mini Kit (QIAGEN 74134). RNA concentration was measured using the Nanodrop OneC, and RNA quality and quantity were validated prior to sequencing. Libraries were titrated using the MiSeq (Illumina) and sequenced on one S4 lane with 2×150bp paired-reads on the Novaseq 6000 (Illumina) at UIUC DNA Services. Sequencing data were processed using the UIUC RNA-seq pipeline, including quality control and read count assessment. Reads were aligned to the Gencode GRCh39 Release 39 (GRCh38.p13) transcriptome. Transcript abundance was quantified using Salmon (v1.5.2) with decoy-aware indexing based on the GRCh38 (primary assembly) genome. Gene-level counts were estimated based on transcript-level counts using tximport. Data are available on NCBI GEO GSE311628.

### Total RNA Sequencing Differential Expression and Pathway Analysis

Differential expression analysis, clustering, and heatmap generation were completed by the Research Informatics Core at UIC. Raw gene-level counts were analyzed using edgeR with differential expression statistics calculated using the exactTest function for pairwise comparisons between experimental groups. P-values were adjusted for multiple testing using the Benjamini-Hochberg false discovery rate (FDR), and genes with an FDR <0.05 were considered significantly differentially expressed. Hierarchical clustering with heatmap construction was performed using the ComplexHeatmap Bioconductor package. The androgen-regulated gene list used for heatmap construction in **Figure 4K** was compiled from nine previously published microarray studies^66^. For pathway-level analysis, gene lists selected from comparisons of interest in differential expression analysis were analyzed through the use of IPA (QIAGEN Inc.) to identify enriched canonical pathways and upstream regulators.

### Quantification and Statistical Analyses

Statistical analyses were performed using Microsoft Excel, Prism v10 (GraphPad), and Seurat (R). Organoid area was compared using non-parametric, unpaired Kruskal-Wallis analysis with Dunn’s post-hoc test to correct for multiple comparisons using statistical hypothesis testing. Mouse body, prostate, or urogenital tissue weights, serum vitamin D hormone levels, and organoid relative area were compared between diets using a parametric, unpaired two-tailed t-test. Gene expression variability was represented as standard deviation in error bars calculated by Prism.

## RESULTS

### Vitamin D enhances prostate epithelial organoid differentiation more than androgen

The mouse prostate has four distinct bilateral lobes, with the dorsal and lateral (DL; dorsolateral) lobes being the most similar to the human peripheral zone by gene expression^32^ (**Figure 1A**); however, no lobes are identical to human.^33^ Cells from each of the lobes were grown into organoids, which expressed both VDR and AR (**Figure 1B-C**). A mixture of solid organoids, translucent organoids, and a small proportion of acinar organoids was observed in organoids from all lobes (**Figure 1C**), which is consistent with previous findings in both mouse organoids^34^ and human organoids.^35^ The organoids differentiated into CK5-expressing basal and CK8/18-expressing luminal epithelial cells (**Figure 1D**). Hormonal crosstalk between vitamin D and androgens is well established in human prostate cancer cell lines.^36–42^ When DL organoids were grown in vehicle control, dihydrotestosterone (DHT), 1,25D, or both DHT+1,25D-supplemented media, the 1,25D and DHT+1,25D conditions resulted in the largest size, indicative of luminal differentiation (**Figure 1E**). These organoids also retained more luminal differentiation capacity after dissociation and reseeding, indicating the preservation of progenitor cell populations (**Figure 1F, Table S1**).

**Figure 1.**
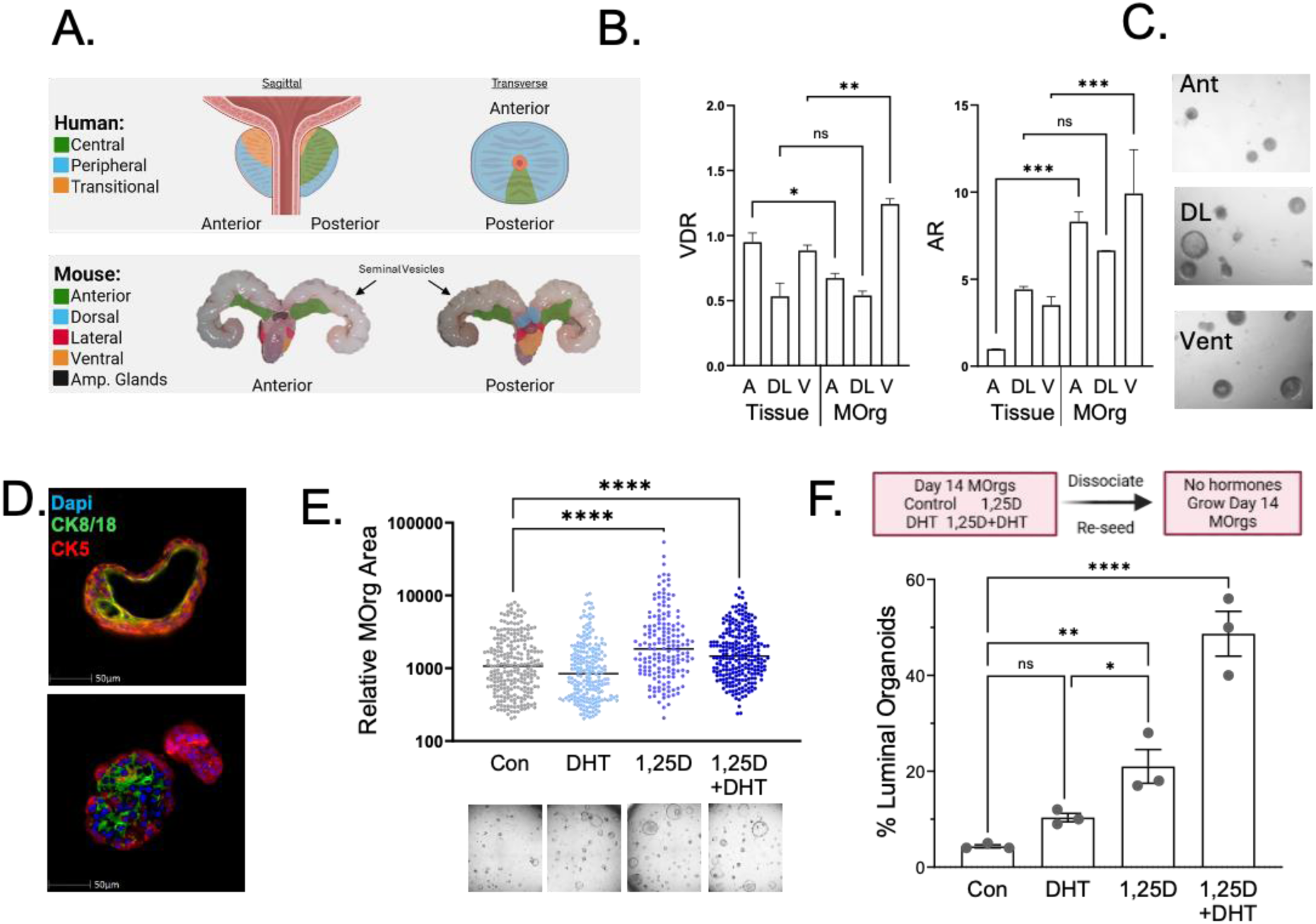
Vitamin D induces differentiation of prostate dorsolateral organoids more than androgens. **A.** Diagram comparing the prostate anatomy between human (zones) and mouse (lobes). **B**, expression of the vitamin D receptor (VDR) and androgen receptor (AR), by RT-qPCR, in mouse prostate anterior (A), dorsolateral (DL) and ventral lobes (V) in whole tissue and day 14 mouse organoids (MOrgs) derived from each lobe. **C**, Representative images of MOrgs from each lobe. **D**, Representative images of luminal (upper panel) and solid (lower panel) DL-MOrgs with both CK8/18 (green) and CK5 (red) positive epithelial populations. **E**, The size of day 14 DL-MOrgs grown in vehicle control (Con), 10 nM 1,25D, 1 nM DHT or both 1,25D+DHT. Representative images are shown below. **F**, The percent of luminal DL-MOrgs after dissociation and reseeding as single cells in the absence of all hormones.

Single-cell RNA sequencing (scRNAseq) of the day 14 DL organoids (**Table S2, Figure S1, GSE304873**) revealed differentiation into five epithelial populations, as well as small populations of fibroblasts and macrophages, as defined by established markers (**Figure 2A-B**). Clear populations were present by cell type, sample, and 1,25D, with the expected expression of key markers in each population (**Figure S2**). The luminal population was the most abundant in organoids grown in 1,25D and in DHT+1,25D (**Figure 2C**). Given the pronounced difference in organoid regeneration (**Figure 1F**), we examined the expression of Tacstd2 (Trop2), which was reported by Goldstein et al^43^ to be responsible for this ability. Organoids grown in 1,25D, both with and without DHT, showed increased Tacstd2 expression in the epithelial clusters (**Figure 2D**). 1,25D-supplementation resulted in many differentially expressed genes (DEGs) in the absence and presence of DHT (**Figure 2E**). These DEGs were significantly enriched in GSEA hallmark pathways related to metabolism, inflammatory signaling, and the estrogen response (**Figure 2F-G**).

**Figure 2.**
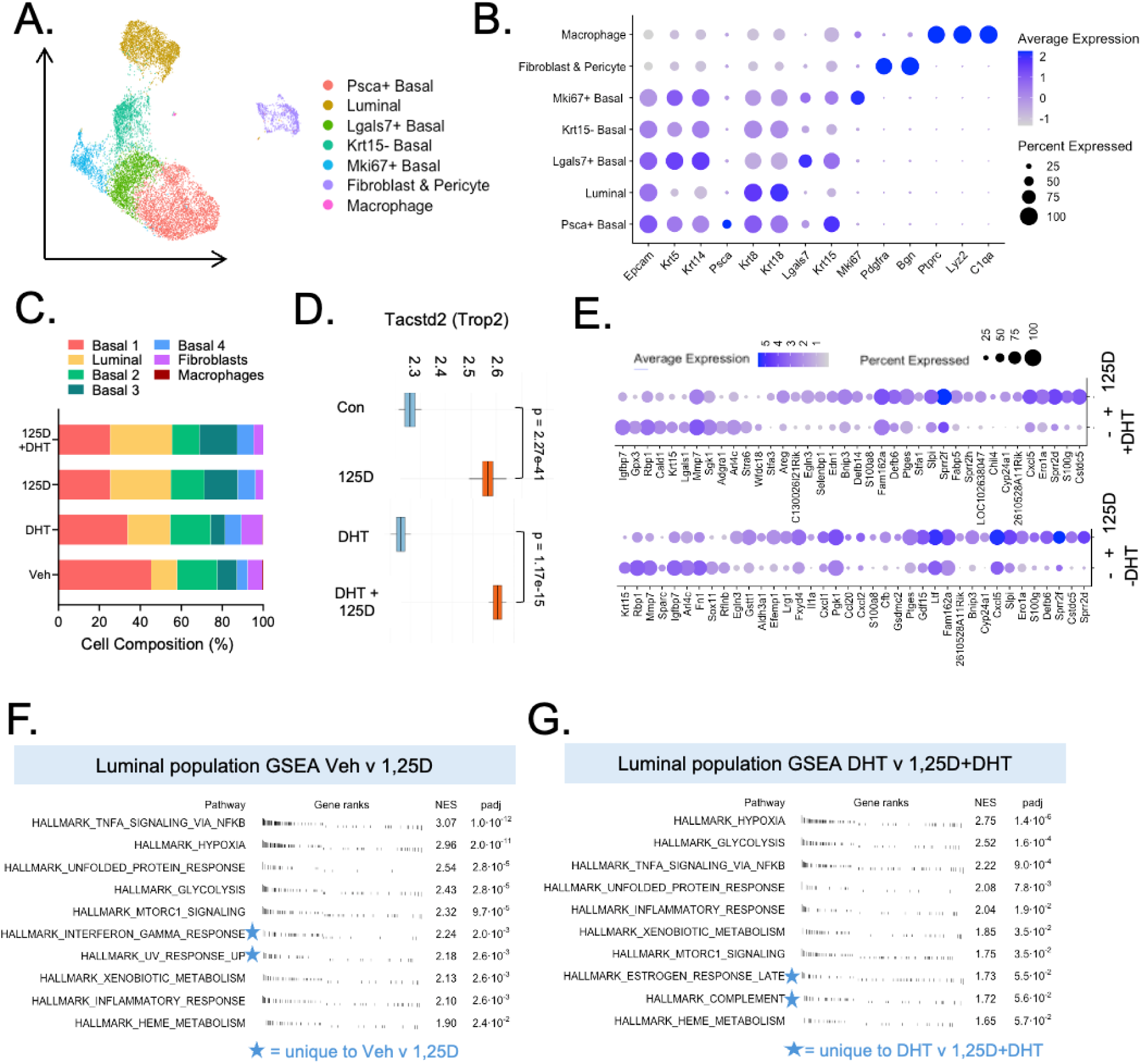
Vitamin D influences luminal cell gene expression and downstream pathways differently in the presence and absence of androgens. **A,** UMAP projection of scRNAseq on day 14 mouse organoids, post-integration. **B.** Dot plot of cell-type markers and differential cluster identifiers for mouse prostate cell populations. **C,** Cluster frequency plot of integrated mouse organoid scRNAseq data displaying the distribution of cells across all 7 clusters between each condition. **D,** Box plots for Tacstd2 (Trop2) expression in epithelial cells from the organoids. **E**, Dotplots of differentially expressed genes (DEGS) in luminal cells between control and 25 nM 1,25D, in the absence and in the presence of 1 nM DHT. DEGS are defined as greater than a 1.5-fold change in expression between conditions and an adjusted p-value < 0.05. **F,** GSEA plot summary tables of top 10 Hallmark pathway genesets comparing luminal cells between control and 25 nM 1,25D, in the absence and **G**, in the presence of 1 nM DHT. Stars indicate unique pathways/genesets comparing vitamin D in the absence or presence of androgens. NES – Normalized Enrichment Score.

### Vitamin D deficiency alters prostate differentiation in vivo

To assess how systemic vitamin D status impacts the prostate in vivo, 8-week-old wild-type male C57BL/6 mice were fed a vitamin D-deficient diet (0.05 IU/g; 50 IU/kg) or a control diet (2.2 IU/g; 2200 IU/kg) for 6 months (**Figure 3A**). The deficient diet reduced serum 25D to below the limit of detection (**Figure 3B**), and these diets have been shown to be safe in rodents with no harmful effects on weight or bone health.^44,45^ We did not observe any effects on mouse weight, prostate size (**Figure S3A**), or the histological appearance of the prostate (**Figure 3C**). Two mice from each diet group were used for scRNAseq (**Table S2, Figure S3, GSE309716**). Previously established cell type markers (cited above) were used for unsupervised Louvain clustering, which identified nine clusters: three epithelial, three stromal, and three immune (**Figure 3D-E**). The epithelial cluster was identified using the expression of Epcam, and the immune cluster was identified using the expression of Ptprc. Within the epithelial cluster, both luminal cells and basal cells were identified. Luminal cells demonstrated expression of Krt8, Krt18, and Nkx3-1, while basal cells demonstrated expression of Krt5, Krt14, and Lgals7 (**Figure 3E**).

**Figure 3.**
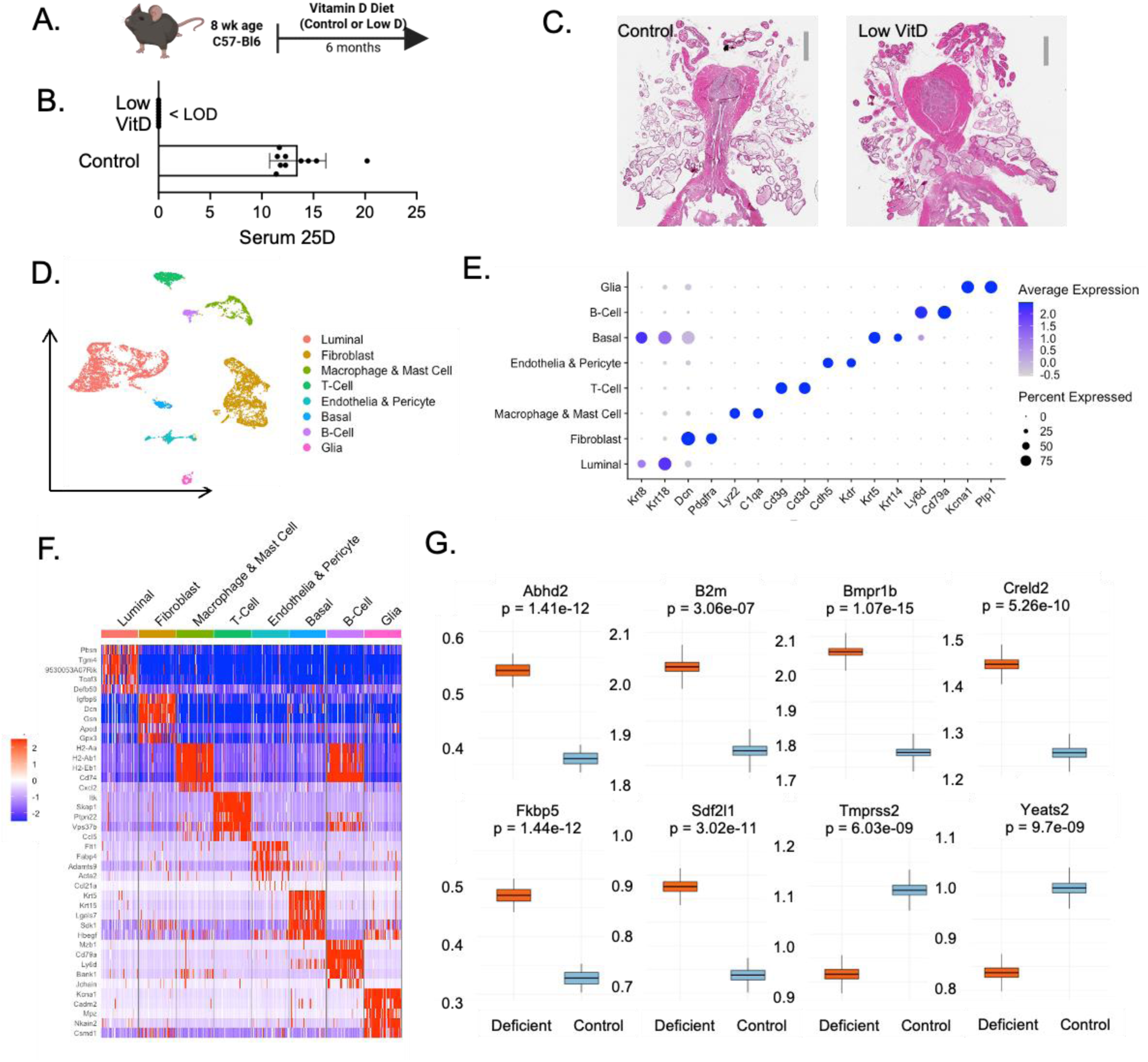
Low vitamin D diet altered gene expression of prostate luminal cells in vivo. **A,** Diagram of the diet study in male mice. **B,** Circulating 25D levels in the serum at the end of the study, LOD was 1.5 ng/mL. **C,** H&E slides of the prostates showed no pathologic abnormalities. **D,** Integrated UMAP dimensionality reduction plot of dorsolateral mouse prostate lobes with unsupervised, Louvain clustering resolution set at 0.05, identifying 8 clusters. In descending size order: luminal cells, fibroblasts, macrophages and mast cells, T-cells, endothelial cells and pericytes, basal cells, B-cells, and glial cells. **E.** Dotplot displaying mouse prostate cell type markers for each cell type present. **F.** Heatmap of the top 5 conserved genes within each cluster sorted by highest average log2 fold change (avg_log2FC) across all samples. Each column represents one cell in each cluster, with 100 cells per cluster. **G.** Box-whisker plots of key DEGS between the diet groups. Y-axis shows Gene Expression mean ± 95% CI.

Using a threshold of adjusted p-value < 0.05, we identified the top 5 DEGs for each cluster (**Figure 3F**). Within the luminal cell population, 37 DEGs were identified, of which 7 were enriched in vitamin D-control luminal cells and 30 were enriched in vitamin D-deficient luminal cells. Most androgen-regulated DEGS, including Fkbp5, Bmpr1b, Sdf2l1, Creld2, B2m, and Abhd2, were upregulated in the deficient diet compared to the sufficient diet, whereas two genes, Yeats2 and Tmprss2, were downregulated (**Figure 3F-G**.). These results reveal the complexity of vitamin D gene regulation and highlight the hormonal crosstalk between the vitamin D and androgen axes, demonstrating the importance of systemic vitamin D status.

### Extended culture in vitamin D-supplement media decreases human prostate cancer cell aggressiveness

To better understand the role of vitamin D in a human cancer model, MDA-PCa2b cells were maintained in vehicle (control), 1 nM or 10 nM 1,25D for six months. This androgen-sensitive cell line was originally derived from a bone metastasis of a 63-year-old African American male and expresses both KLK3/PSA and AR^39,46,47^ as well as functional VDR,^39^ making them an ideal model to study the hormonal crosstalk between vitamin D and androgens. The 1,25D-adapted MDA-PCa-2b cells were grown as xenografts in SCID mice (n=10 grafts/treatment). Between 18 and 25 weeks post-injection, each xenograft derived from the vehicle control cells had formed a palpable, measurable, PSA-producing tumor, with all tumors reaching 500 mm³ by 175 days (**Figure 4B-C**). The 1 nM and 10 nM 1,25D-treated MDA-PCa-2b cells did not grow tumors and did not have detectable serum PSA (**Figure 4B-C**). By contrast, in vitro sustained 1,25D treatment of MDA-PCa-2b cells for 6 months resulted in a slightly higher proliferation rate than acute 1,25D treatment for only 96 hours (**Figure 4D**).

**Figure 4.**
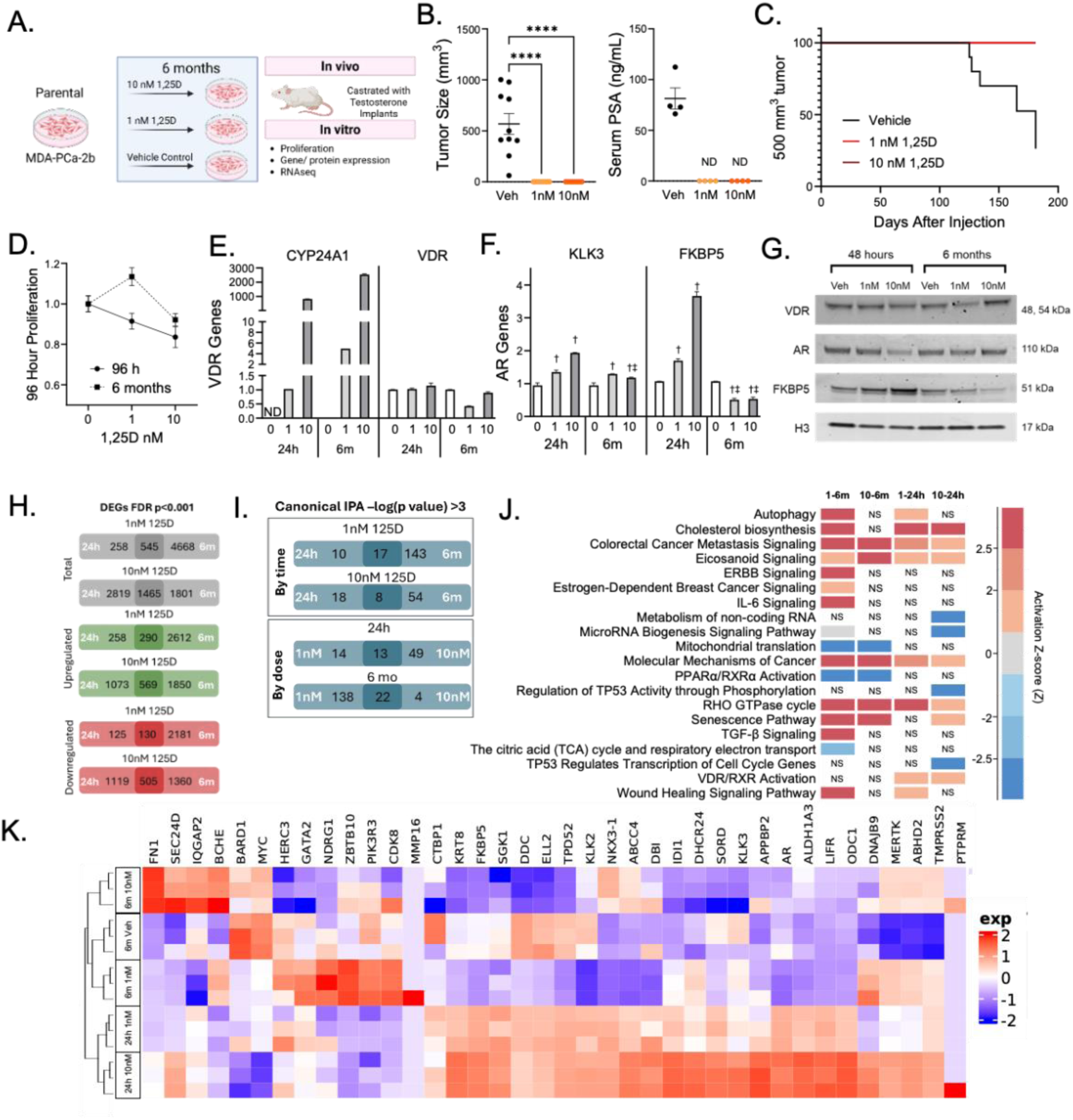
MDA-PCa-2b prostate cancer cells, serially passaged in 1,25D, exhibit diminished xenograft formation and altered androgen signaling. **A,** Study design diagram. MDA-PCa-2b cells were maintained in media containing 1 nM 1,25D, 10 nM 1,25D, or vehicle control for 6 months to create adapted cells. Adapted cells (10 grafts per cell line) were injected into castrated SCID mice with T-implants. **B**, Tumor size and serum PSA levels at endpoint/death for each cell line. Kruskal-Wallis with Dunn’s multiple comparisons test t (****p < 0.0001). **C,** Kaplan-Meier survival analysis with the event of reaching a 500 mm³-sized tumor. **D,** In vitro proliferation at 96 hours of the 6-month treated cells compared to control cells treated for 96 hours with 1 and 10 nM 1,25D. **E,** Expression of vitamin D-regulated genes, and F, androgen-regulated genes, in the 6-month treated cells compared to control cells treated for 24 hours with 1 and 10 nM 1,25D. **G,** Protein levels of VDR, AR, and FKBP5 in the 6-month treated cells compared to control cells treated for 48 hours with 1 and 10 nM 1,25D. **H,** Summary of DEGS, and **I,** Ingenuity Pathway Analyses (IPA) by bulk RNAseq between the cell lines. **J,** comparison of top canonical IPAs between the cell lines with p<0.01. **K,** The expression of androgen pathway genes in RNAseq from the 6-month treated cells compared to control cells treated for 24 hours with 1 and 10 nM 1,25D.

The extended culture of MDA-PCa-2b cells in 1,25D resulted in a biphasic suppression of VDR at the 1 nM dose, accompanied by the expected increase in CYP24A1, a robustly 1,25D-upregulated gene (**Figure 4E**). Crosstalk with the androgen pathway differed by the duration of 1,25D treatment, with upregulation at 24h while, extended 6-month treatment showed attenuated KLK3 expression and suppression of FKBP5 (**Figure 4F**). FKBP5 protein was also reduced in the extended cultures, despite no difference in AR protein levels (**Figure 4G**).

Bulk RNA-seq was performed in triplicate on the cells and compared to 24-hour culture in 1 and 10 nM 1,25D (the 6-month vehicle-treated cells were used as controls for the 24-hour 1,25D treatments) (GSE311628). The 6-month 1 nM 1,25D dose had the highest number of DEGs (5213), with only 11.6% overlap with the DEGs from the 24h treatment at 1nM (**Figure 4H**). Ingenuity Pathway Analysis (IPA) showed common and unique pathways among the treatments (**Figure 4I**, **Table S4**). Pathways with the highest z-scores showed treatment-dependent variability, demonstrating distinct transcriptional regulatory patterns in cells maintained in vitamin D for 6 months (**Figure 4J**). Analyses of previously established AR-regulated genes showed opposing effects between 24-hour and 6-month treatments for many of the genes (**Figure 4K**).

## DISCUSSION

Prior prostate organoid studies have consistently demonstrated that androgens are required for luminal epithelial differentiation, regardless of the epithelial stem or progenitor cell of origin.^43,48,49^ For example, Shen et al. showed that DHT supplementation promotes the formation of larger, more differentiated luminal organoids compared with androgen-deprived conditions. Consistent with these findings, we observed enhanced luminal differentiation with DHT supplementation. However, our data revealed an additional layer of hormonal regulation in which vitamin D, either alone or in combination with DHT, promotes luminal epithelial differentiation to a greater extent than DHT alone. These findings extend our previous work in human prostate epithelial organoids, demonstrating accelerated luminal differentiation following vitamin D treatment.^50^ ScRNAseq further identified unique pathways, including late estrogen response and complement signaling, in luminal cells that are specifically activated by vitamin D or DHT supplementation. Notably, Tacsdt2 expression was increased in the epithelial organoids, consistent with findings by Goldstein et al, who found Tacsdt2 (Trop2) was an essential protein for prostate organoid regeneration^43^. The use of mouse-derived organoids enabled a controlled assessment of vitamin D signaling in a genetically and environmentally uniform system, thereby strengthening the biological relevance of these observations.

Given the high prevalence of vitamin D deficiency in populations disproportionately affected by aggressive prostate cancer, there is a possibility that chronic vitamin D deficiency throughout an individual’s lifespan may alter the prostate epithelium toward a state permissive for more aggressive disease. Other vitamin D-deficiency diet studies have focused on PCa growth and metastasis in various genetically engineered mouse models and human PCa xenografts.^23,24,51^ One study reported increased prostatic hyperplasia in middle-aged mice maintained on a vitamin D-deficient diet from birth, which was partially reversed upon repletion with a vitamin D-sufficient diet.^52^ Our in vivo model isolates the effect of dietary vitamin D deficiency on the normal prostate, allowing us to define how vitamin D status alone shapes prostate epithelial biology. At the single-cell level, mice maintained on a vitamin D-deficient diet exhibited marked changes in androgen-responsive gene expression in luminal epithelial cells, despite the absence of gross or histologic prostate abnormalities. Notably, elevated expression of Fkbp5 and loss of Tmprss2 were observed, two established functional indicators of androgen signaling, pointing to a transcriptional reprogramming of androgen responsiveness in prostate luminal cells by vitamin D deficiency. Such transcriptional changes arising solely from reduced vitamin D status may represent early molecular events that precede disease initiation or progression, consistent with epidemiologic observations linking vitamin D deficiency to aggressive, rather than indolent, prostate cancer.

A key novelty of our MDA-PCa-2b experiments is the distinction between short-term exposure and long-term hormonal adaptation. In vitro studies typically assess treatment effects using short-term, pulse-based exposures across varying doses; however, such approaches do n ot capture the consequences of sustained hormonal signaling that the cells experience in situ. To address this, we examined the effects of two physiologically relevant doses of 1,25D under both acute treatment and long-term 6-month adaptation. This long-term adaptation induced extensive transcriptional reprogramming that was distinct from short-term treatment effects. For both doses of 1,25D, the majority of differentially expressed genes were unique to the adapted cells, with only modest overlap with those in acutely treated cells, indicating a durable shift in the cellular state rather than a transient response. Acute 1,25D treatment largely preserved or enhanced a canonical androgen-driven program, marked by increased expression of AR target genes and luminal differentiation markers, including KLK2, KLK3, FKBP5, NXK3-1, SGK1, and KRT8, compared with control or long-term-treated cells. In contrast, long-term exposure to 1,25D induced distinct dose-dependent transcriptional programs: low-dose adaptation (1nM) was characterized by enrichment of transcriptional regulators and signaling modulators such as GATA2, NDRG1, and CDK8, whereas high-dose adaptation (10nM) uniquely induced genes associated with extracellular matrix organization, cytoskeleton stability, and cellular anchoring, including FN1 and IQGAP2. Consistent with this notion, canonical pathway analysis revealed numerous pathways uniquely enriched in long-term adapted cells. By contrast, the canonical VDR-RXR activation pathway was enriched only in cells treated for 24 hours, and not in 6-month adapted cells, further highlighting the bias in existing databases toward shorter hormone treatment pulses. Functionally, 1,25D-adapted cells failed to form tumors upon implantation of xenografted PCa cells, importantly demonstrating that chronic vitamin D exposure alters tumorigenic potential. Together, these findings highlight the contrasting phenotypes of pulse hormone treatment models and chronically treated cell models, demonstrating that sustained vitamin D signaling reprograms PCa cells toward a less aggressive or even indolent state.

Our findings and approach fill a gap between the epidemiological links between vitamin D and PCa and interventional trials with vitamin D supplementation. There are significant challenges to prospective interventional trials with vitamin D supplementation that include adherence, inadequate duration of the intervention, and low incidence of aggressive-lethal PCa during the study period. Dietary deficiency animal models mimic real-life conditions in patients, but in a more controlled environment allowing distinction of vitamin D effects from other variables.

Importantly, this study highlights that vitamin D deficiency alters prostate biology even in the absence of overt disease. Changes in epithelial differentiation and androgen-responsive transcriptional programs were observed prior to tumor formation, suggesting that vitamin D status may influence prostate cancer risk by shaping the cellular and molecular landscapes of the prostate over time. These data support the use of a model in which vitamin D sufficiency promotes epithelial differentiation and hormonal balance, while deficiency creates a permissive environment for the development and progression of benign and malignant disease. Our study does not address excess vitamin D exposure but instead focuses on biologically relevant deficiency, which disproportionately affects non-Hispanic Black men in the U.S. Given the higher PCa mortality and higher prevalence of vitamin D deficiency in these men compared to other populations, our findings provide mechanistic support for the hypothesis that vitamin D deficiency contributes to their greater likelihood of developing aggressive PCa.

In summary, our work demonstrates that vitamin D is a critical regulator of prostatic epithelial differentiation, androgen signaling, and of the risk of developing aggressive prostate cancer across multiple biologically relevant models of human disease. These findings reinforce the notion of vitamin D as an essential hormone in prostate biology and provide a mechanistic framework for understanding its protective role against aggressive prostate cancer. Defining optimal vitamin D status and understanding its long-term effects on prostate biology and health may represent an important strategy for reducing prostate cancer disparities and improving outcomes in vulnerable populations.

## CONTRIBUTIONS OF AUTHORS

Conceptualization: L.N., K.D.K., R.M.S. Data Acquisition: K.D.K., R.A.H., S.C., R.M.B., L.W., A.D., M.J.S. Formal Analysis: K.D.K., M.C.B., R.A.H., S.C., A.D. Manuscript Preparation: L.N., A.D., K.D.K., M.C.B., R.M.S., L.W. Supervision: L.N., D.V.G., R.M.S. Funding Acquisition: L.N., R.M.S.

## ACKNOWLEDGEMENTS

Next-generation sequencing and sequence alignment were completed by the University of Illinois at the Urbana-Champaign Roy J. Carver Biotechnology Center, Drs. Jenny D. Zadeh and Alvaro G. Hernandez. Animal care was at the University of Illinois Chicago (UIC) Animal Care Facility. Bioinformatics analysis for the bulkRNAseq was provided by Leonid Feferman and Mark Maienschein-Cline in the UIC Research Informatics Core (RIC), supported in part by NCATS through Grant UM1TR005438, and the UIC Cancer Center Cancer Bioinformatics Shared Resource (CBSR). Some figures were created using BioRender.com.

## FUNDING

University of Illinois Cancer Center Pilot Project Award (Nonn and Sargis).

Department of Defense USAMRAA W81XWH2010182 (Nonn).

National Institute of Health R01CA290811 (Nonn).

