## Supplemental Tables 1-4 and Supplemental Figures 1-3 for "Vitamin D deficiency alters prostate epithelial differentiation and increases prostate cancer aggressiveness in ex vivo and in vivo models"

### Slide 1
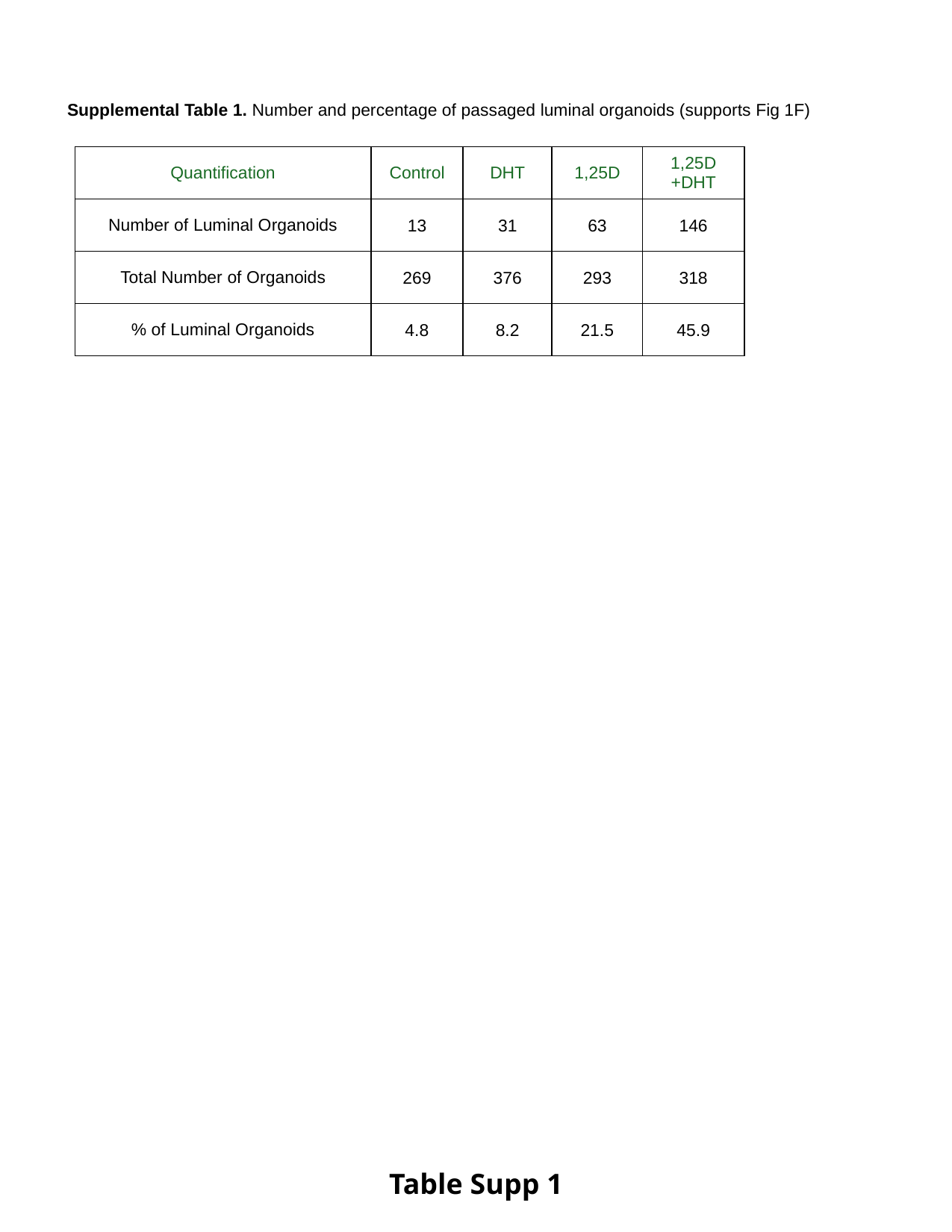

Supplemental Table 1. Number and percentage of passaged luminal organoids (supports Fig 1F)
| Quantification | Control | DHT | 1,25D | 1,25D +DHT |
| --- | --- | --- | --- | --- |
| Number of Luminal Organoids | 13 | 31 | 63 | 146 |
| Total Number of Organoids | 269 | 376 | 293 | 318 |
| % of Luminal Organoids | 4.8 | 8.2 | 21.5 | 45.9 |
Table Supp 1

### Slide 2
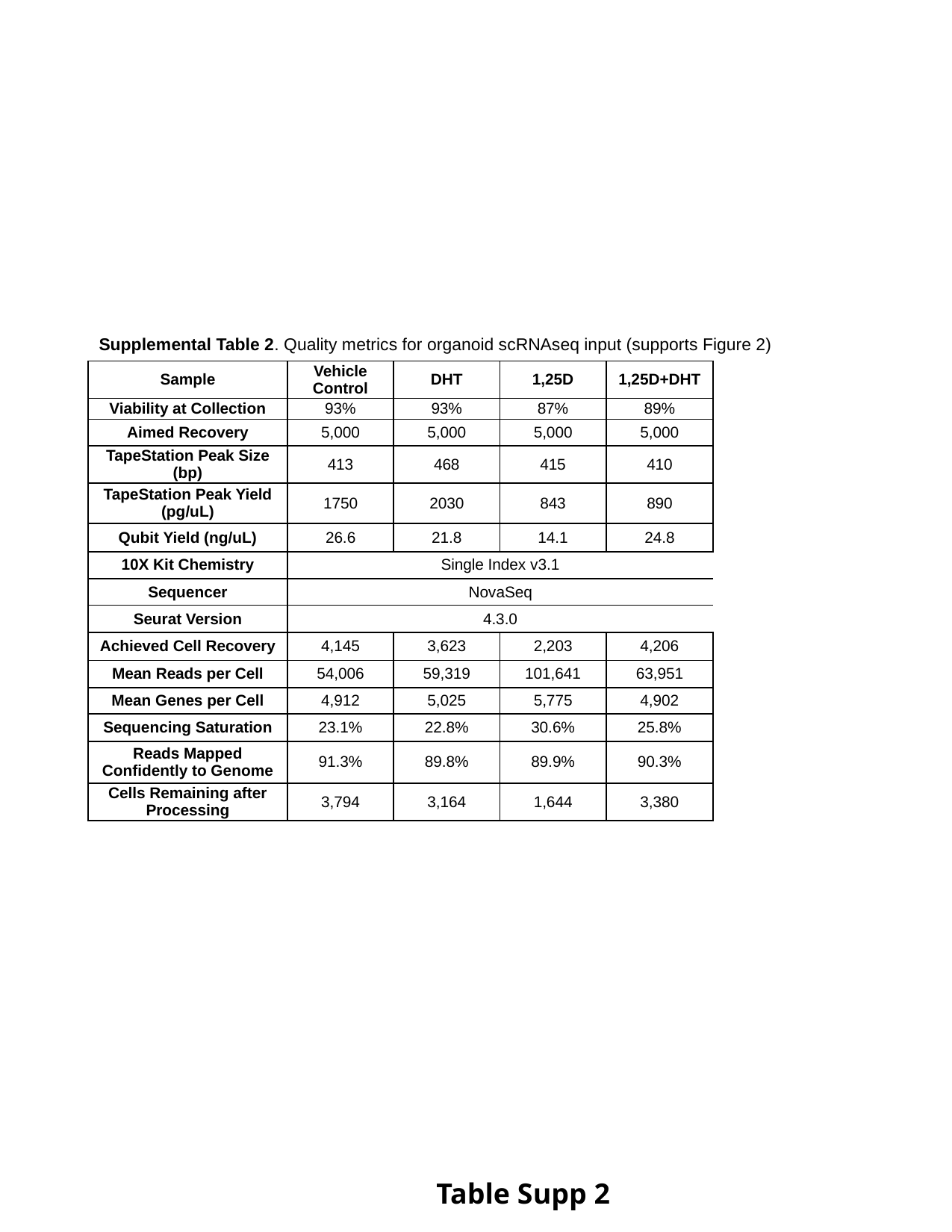

Supplemental Table 2. Quality metrics for organoid scRNAseq input (supports Figure 2)
| Sample | Vehicle Control | DHT | 1,25D | 1,25D+DHT |
| --- | --- | --- | --- | --- |
| Viability at Collection | 93% | 93% | 87% | 89% |
| Aimed Recovery | 5,000 | 5,000 | 5,000 | 5,000 |
| TapeStation Peak Size (bp) | 413 | 468 | 415 | 410 |
| TapeStation Peak Yield (pg/uL) | 1750 | 2030 | 843 | 890 |
| Qubit Yield (ng/uL) | 26.6 | 21.8 | 14.1 | 24.8 |
| 10X Kit Chemistry | Single Index v3.1 | | | |
| Sequencer | NovaSeq | | | |
| Seurat Version | 4.3.0 | | | |
| Achieved Cell Recovery | 4,145 | 3,623 | 2,203 | 4,206 |
| Mean Reads per Cell | 54,006 | 59,319 | 101,641 | 63,951 |
| Mean Genes per Cell | 4,912 | 5,025 | 5,775 | 4,902 |
| Sequencing Saturation | 23.1% | 22.8% | 30.6% | 25.8% |
| Reads Mapped Confidently to Genome | 91.3% | 89.8% | 89.9% | 90.3% |
| Cells Remaining after Processing | 3,794 | 3,164 | 1,644 | 3,380 |
Table Supp 2

### Slide 3
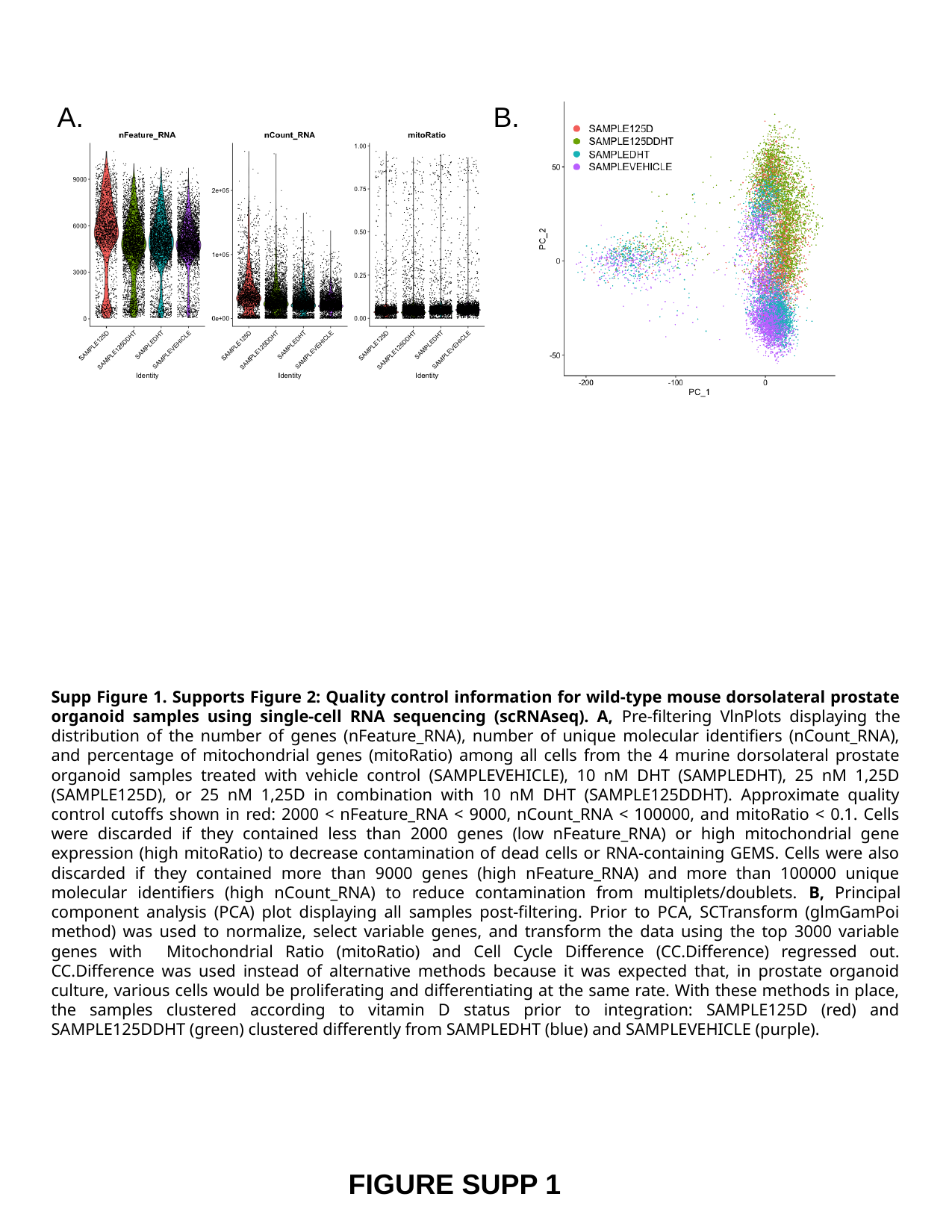

A.
B.
Supp Figure 1. Supports Figure 2: Quality control information for wild-type mouse dorsolateral prostate organoid samples using single-cell RNA sequencing (scRNAseq). A, Pre-filtering VlnPlots displaying the distribution of the number of genes (nFeature_RNA), number of unique molecular identifiers (nCount_RNA), and percentage of mitochondrial genes (mitoRatio) among all cells from the 4 murine dorsolateral prostate organoid samples treated with vehicle control (SAMPLEVEHICLE), 10 nM DHT (SAMPLEDHT), 25 nM 1,25D (SAMPLE125D), or 25 nM 1,25D in combination with 10 nM DHT (SAMPLE125DDHT). Approximate quality control cutoffs shown in red: 2000 < nFeature_RNA < 9000, nCount_RNA < 100000, and mitoRatio < 0.1. Cells were discarded if they contained less than 2000 genes (low nFeature_RNA) or high mitochondrial gene expression (high mitoRatio) to decrease contamination of dead cells or RNA-containing GEMS. Cells were also discarded if they contained more than 9000 genes (high nFeature_RNA) and more than 100000 unique molecular identifiers (high nCount_RNA) to reduce contamination from multiplets/doublets. B, Principal component analysis (PCA) plot displaying all samples post-filtering. Prior to PCA, SCTransform (glmGamPoi method) was used to normalize, select variable genes, and transform the data using the top 3000 variable genes with Mitochondrial Ratio (mitoRatio) and Cell Cycle Difference (CC.Difference) regressed out. CC.Difference was used instead of alternative methods because it was expected that, in prostate organoid culture, various cells would be proliferating and differentiating at the same rate. With these methods in place, the samples clustered according to vitamin D status prior to integration: SAMPLE125D (red) and SAMPLE125DDHT (green) clustered differently from SAMPLEDHT (blue) and SAMPLEVEHICLE (purple).
FIGURE SUPP 1

### Slide 4
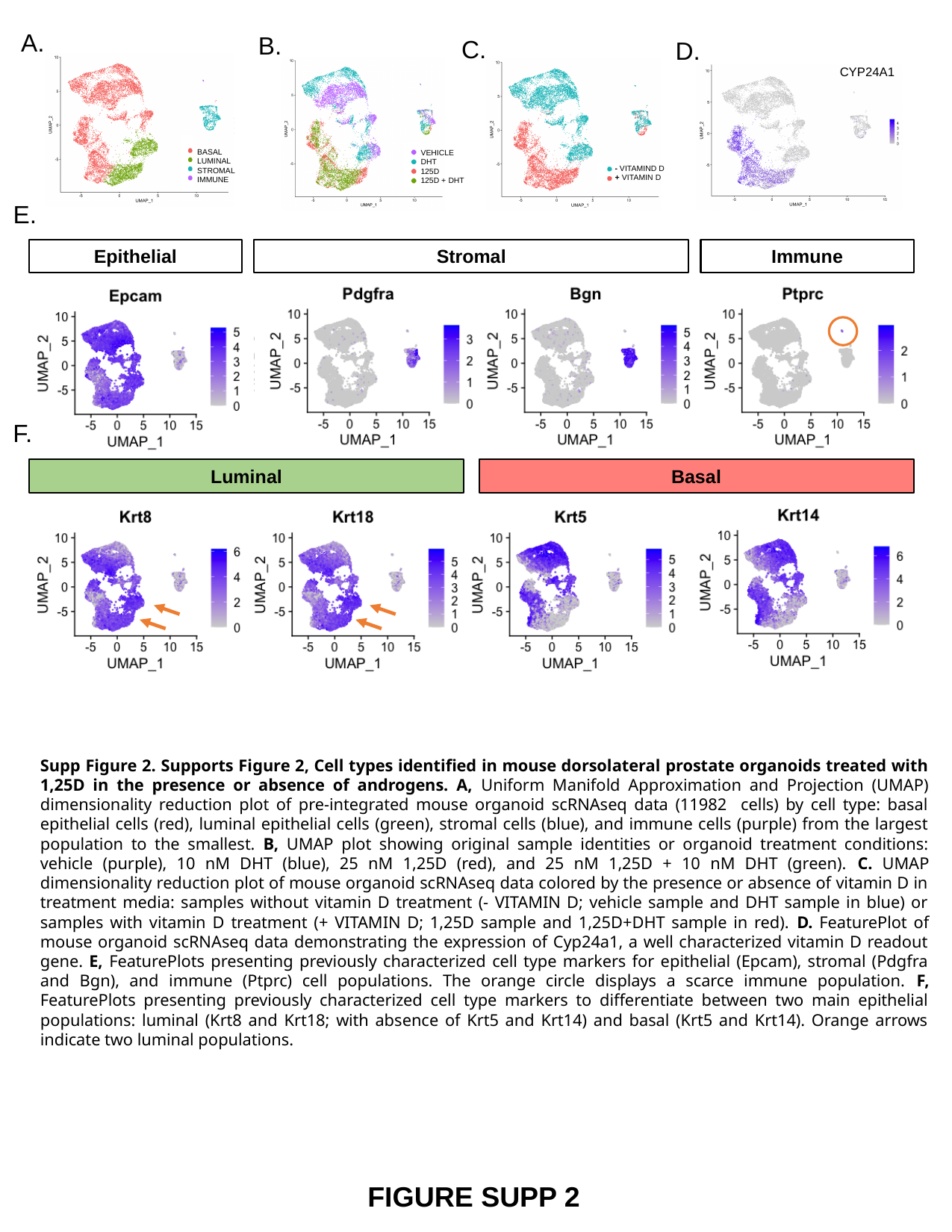

A.
B.
C.
D.
CYP24A1
BASAL
LUMINAL
STROMAL
IMMUNE
VEHICLE
DHT
125D
125D + DHT
- VITAMIND D
+ VITAMIN D
E.
Epithelial
Stromal
Immune
F.
Luminal
Basal
Supp Figure 2. Supports Figure 2, Cell types identified in mouse dorsolateral prostate organoids treated with 1,25D in the presence or absence of androgens. A, Uniform Manifold Approximation and Projection (UMAP) dimensionality reduction plot of pre-integrated mouse organoid scRNAseq data (11982 cells) by cell type: basal epithelial cells (red), luminal epithelial cells (green), stromal cells (blue), and immune cells (purple) from the largest population to the smallest. B, UMAP plot showing original sample identities or organoid treatment conditions: vehicle (purple), 10 nM DHT (blue), 25 nM 1,25D (red), and 25 nM 1,25D + 10 nM DHT (green). C. UMAP dimensionality reduction plot of mouse organoid scRNAseq data colored by the presence or absence of vitamin D in treatment media: samples without vitamin D treatment (- VITAMIN D; vehicle sample and DHT sample in blue) or samples with vitamin D treatment (+ VITAMIN D; 1,25D sample and 1,25D+DHT sample in red). D. FeaturePlot of mouse organoid scRNAseq data demonstrating the expression of Cyp24a1, a well characterized vitamin D readout gene. E, FeaturePlots presenting previously characterized cell type markers for epithelial (Epcam), stromal (Pdgfra and Bgn), and immune (Ptprc) cell populations. The orange circle displays a scarce immune population. F, FeaturePlots presenting previously characterized cell type markers to differentiate between two main epithelial populations: luminal (Krt8 and Krt18; with absence of Krt5 and Krt14) and basal (Krt5 and Krt14). Orange arrows indicate two luminal populations.
FIGURE SUPP 2

### Slide 5
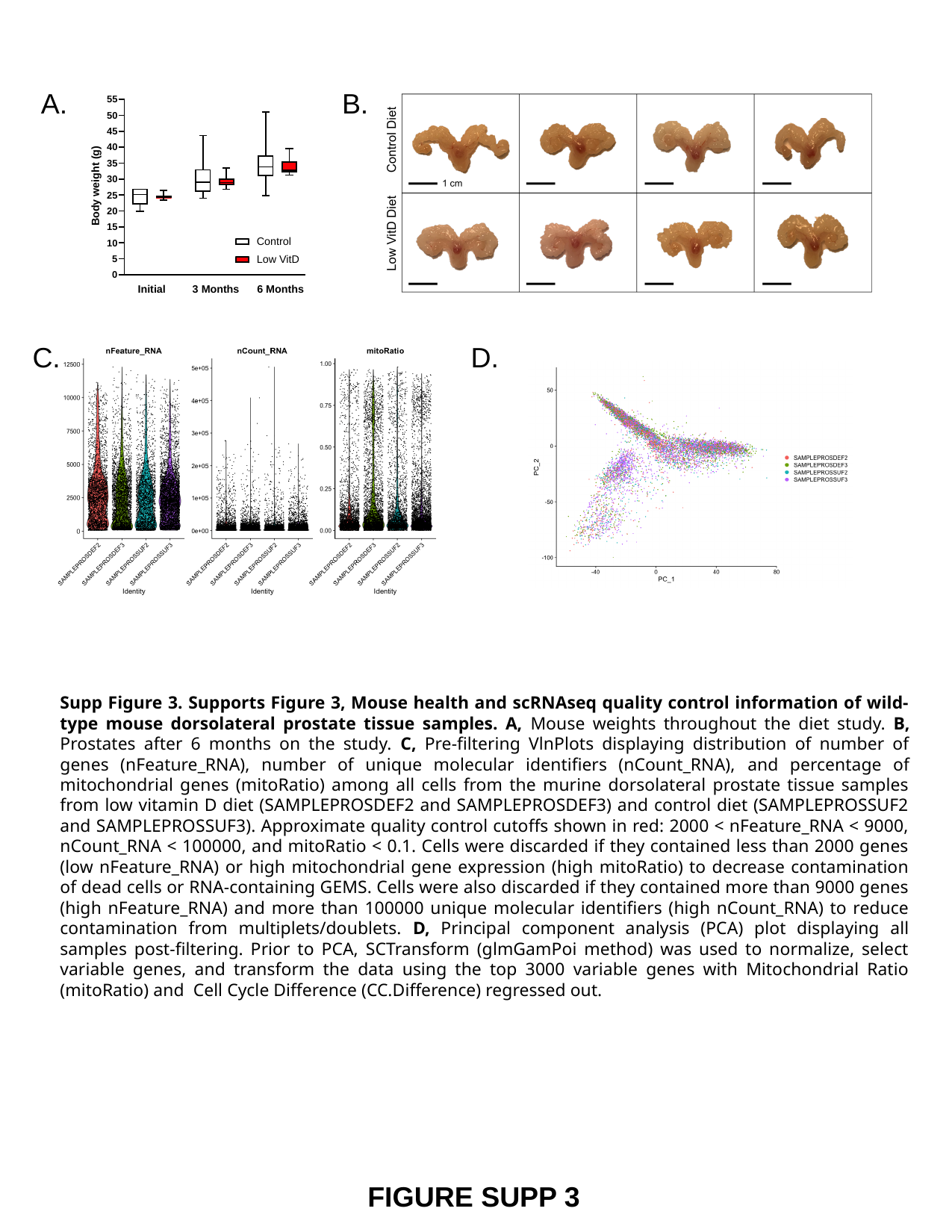

A.
B.
C.
D.
Supp Figure 3. Supports Figure 3, Mouse health and scRNAseq quality control information of wild-type mouse dorsolateral prostate tissue samples. A, Mouse weights throughout the diet study. B, Prostates after 6 months on the study. C, Pre-filtering VlnPlots displaying distribution of number of genes (nFeature_RNA), number of unique molecular identifiers (nCount_RNA), and percentage of mitochondrial genes (mitoRatio) among all cells from the murine dorsolateral prostate tissue samples from low vitamin D diet (SAMPLEPROSDEF2 and SAMPLEPROSDEF3) and control diet (SAMPLEPROSSUF2 and SAMPLEPROSSUF3). Approximate quality control cutoffs shown in red: 2000 < nFeature_RNA < 9000, nCount_RNA < 100000, and mitoRatio < 0.1. Cells were discarded if they contained less than 2000 genes (low nFeature_RNA) or high mitochondrial gene expression (high mitoRatio) to decrease contamination of dead cells or RNA-containing GEMS. Cells were also discarded if they contained more than 9000 genes (high nFeature_RNA) and more than 100000 unique molecular identifiers (high nCount_RNA) to reduce contamination from multiplets/doublets. D, Principal component analysis (PCA) plot displaying all samples post-filtering. Prior to PCA, SCTransform (glmGamPoi method) was used to normalize, select variable genes, and transform the data using the top 3000 variable genes with Mitochondrial Ratio (mitoRatio) and Cell Cycle Difference (CC.Difference) regressed out.
FIGURE SUPP 3

### Slide 6
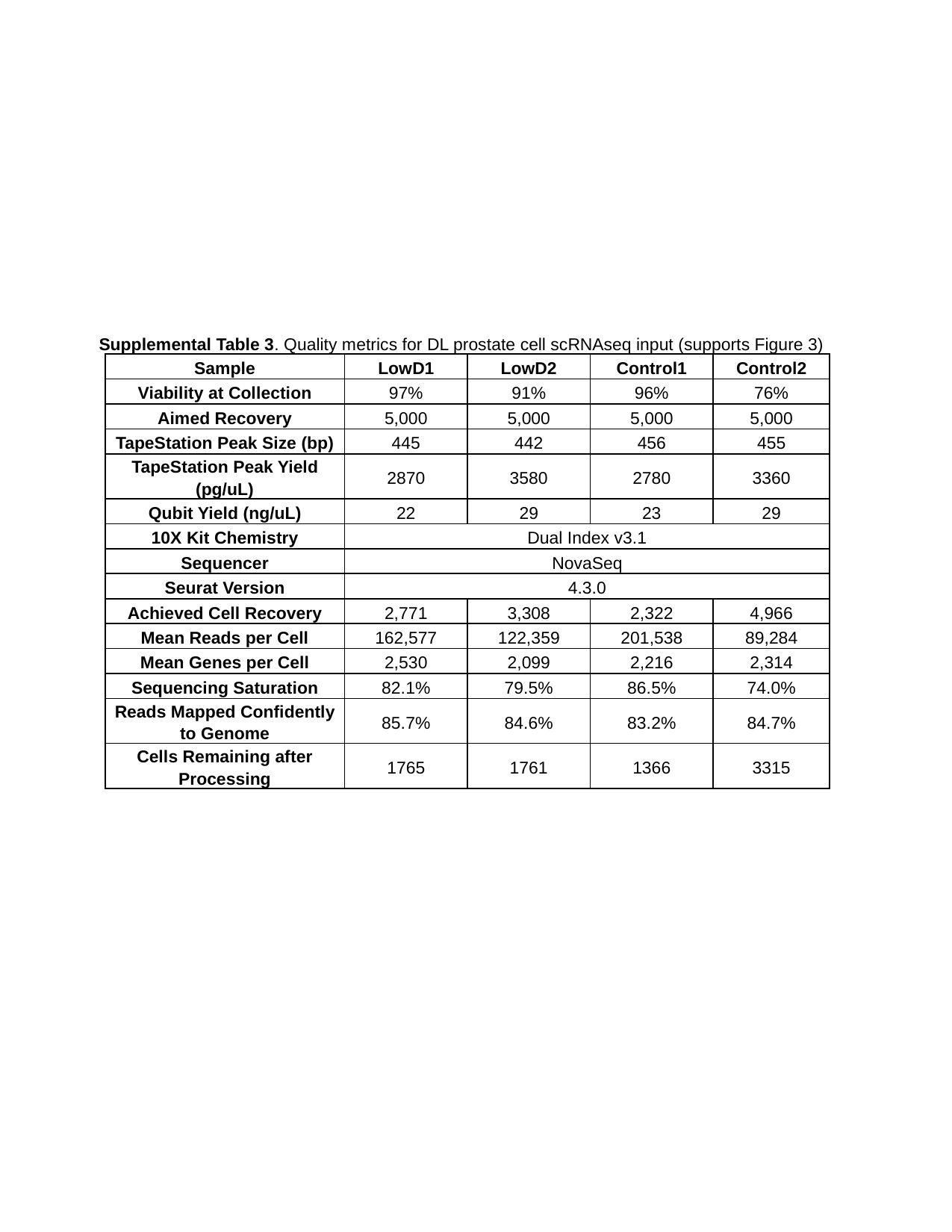

Supplemental Table 3. Quality metrics for DL prostate cell scRNAseq input (supports Figure 3)
| Sample | LowD1 | LowD2 | Control1 | Control2 |
| --- | --- | --- | --- | --- |
| Viability at Collection | 97% | 91% | 96% | 76% |
| Aimed Recovery | 5,000 | 5,000 | 5,000 | 5,000 |
| TapeStation Peak Size (bp) | 445 | 442 | 456 | 455 |
| TapeStation Peak Yield (pg/uL) | 2870 | 3580 | 2780 | 3360 |
| Qubit Yield (ng/uL) | 22 | 29 | 23 | 29 |
| 10X Kit Chemistry | Dual Index v3.1 | | | |
| Sequencer | NovaSeq | | | |
| Seurat Version | 4.3.0 | | | |
| Achieved Cell Recovery | 2,771 | 3,308 | 2,322 | 4,966 |
| Mean Reads per Cell | 162,577 | 122,359 | 201,538 | 89,284 |
| Mean Genes per Cell | 2,530 | 2,099 | 2,216 | 2,314 |
| Sequencing Saturation | 82.1% | 79.5% | 86.5% | 74.0% |
| Reads Mapped Confidently to Genome | 85.7% | 84.6% | 83.2% | 84.7% |
| Cells Remaining after Processing | 1765 | 1761 | 1366 | 3315 |

### Slide 7
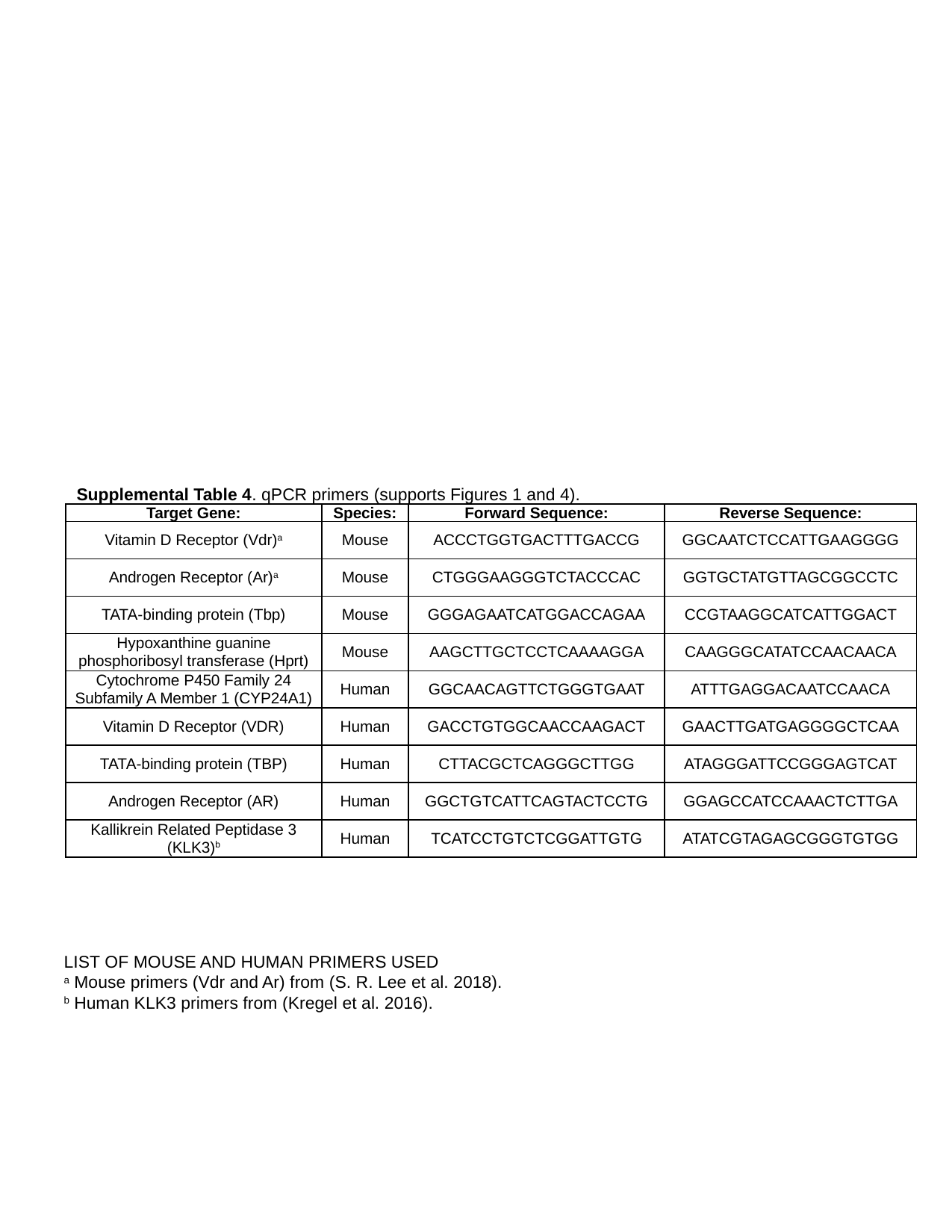

Supplemental Table 4. qPCR primers (supports Figures 1 and 4).
| Target Gene: | Species: | Forward Sequence: | Reverse Sequence: |
| --- | --- | --- | --- |
| Vitamin D Receptor (Vdr)^a^ | Mouse | ACCCTGGTGACTTTGACCG | GGCAATCTCCATTGAAGGGG |
| Androgen Receptor (Ar)^a^ | Mouse | CTGGGAAGGGTCTACCCAC | GGTGCTATGTTAGCGGCCTC |
| TATA-binding protein (Tbp) | Mouse | GGGAGAATCATGGACCAGAA | CCGTAAGGCATCATTGGACT |
| Hypoxanthine guanine phosphoribosyl transferase (Hprt) | Mouse | AAGCTTGCTCCTCAAAAGGA | CAAGGGCATATCCAACAACA |
| Cytochrome P450 Family 24 Subfamily A Member 1 (CYP24A1) | Human | GGCAACAGTTCTGGGTGAAT | ATTTGAGGACAATCCAACA |
| Vitamin D Receptor (VDR) | Human | GACCTGTGGCAACCAAGACT | GAACTTGATGAGGGGCTCAA |
| TATA-binding protein (TBP) | Human | CTTACGCTCAGGGCTTGG | ATAGGGATTCCGGGAGTCAT |
| Androgen Receptor (AR) | Human | GGCTGTCATTCAGTACTCCTG | GGAGCCATCCAAACTCTTGA |
| Kallikrein Related Peptidase 3 (KLK3)^b^ | Human | TCATCCTGTCTCGGATTGTG | ATATCGTAGAGCGGGTGTGG |
LIST OF MOUSE AND HUMAN PRIMERS USED
^a^ Mouse primers (Vdr and Ar) from (S. R. Lee et al. 2018).
^b^ Human KLK3 primers from (Kregel et al. 2016).
